# Visual Field Inhomogeneities and the Architectonics of Early Visual Cortex Shape Visual Working Memory

**DOI:** 10.1101/2025.05.13.653695

**Authors:** Julia Papiernik-Kłodzińska, Simon Hviid Del Pin, Kristian Sandberg, Michał Wierzchoń, Marisa Carrasco, Renate Rutiku

## Abstract

Whether the primary visual cortex (V1) is essential for visual working memory (vWM) remains a topic of scientific debate. The current study expanded upon previous findings by examining whether idiosyncratic architectural properties of V1, particularly those underlying visual field inhomogeneities such as polar angle asymmetries, predict interindividual differences in vWM performance. A total of 292 participants underwent quantitative MRI (qMRI) using a multiparametric mapping sequence to generate four microstructural maps per participant: magnetization transfer (MT), proton density (PD), longitudinal relaxation rate (R1), and transverse relaxation rate (R2*). In a separate session, the participants completed a vWM task designed to probe visual field inhomogeneities. Behavioral results are consistent with previously reported asymmetries in vWM, particularly the inverted polarity of the vertical meridian asymmetry (VMA). Quantitative MRI analysis revealed significant associations between VMA and multiple qMRI parameters within V1, indicating that V1 microarchitecture contributes to variability in vWM performance. Additionally, cortical thickness measures linked V3 to left–right asymmetry, suggesting that structural variability in the early visual cortex beyond V1 also shapes vWM performance. These findings are consistent with the sensory recruitment hypothesis and demonstrate that fine-grained architectural characteristics of early visual areas constrain vWM performance.

---

Working memory is the core cognitive ability to “hold a limited amount of information temporarily in a heightened state of availability for use in ongoing information processing” (Cowan 2017). Consequently, the term ‘visual working memory’ (vWM) is used to specifically denote the temporary storage and use of visually perceived information (Lorenc et al. 2021). It is known that vWM is severely limited in capacity, and interindividual differences in capacity limits are strongly linked to other cognitive abilities, such as intelligence (Fukuda et al. 2010). For this reason, there is considerable interest in why people with otherwise comparable perceptual abilities can differ so markedly in their ability to retain perceived information over time (Luck and Vogel 2013). The two hallmarks of vWM – its constrained capacity and stable interindividual differences – have been instrumental in guiding the search for its neural underpinnings.

Due to the limited capacity of working memory, it was originally thought that separate modality-specific storage exists independently of high-capacity sensory cortices with their rich perceptual representations. It was theorized that perceptual contents are first encoded into more abstract representations and subsequently retained in frontal and/or parietal cortices. Indeed, the landmark monkey study by Funahashi and colleagues (Funahashi et al. 1989) delivered compelling evidence for such a model (see also Miller et al. 1996). However, this separation between perception and vWM retention was challenged by Harrison & Tong (Harrison and Tong 2009) and Serences and colleagues (Serences et al. 2009), who demonstrated that vWM content can be tracked within the primary visual cortex (V1). Many studies followed with similar findings (see Ma et al. 2014 and Christophel et al. 2017 for reviews), leading to the sensory recruitment hypothesis. This model suggests that early sensory areas are involved in maintaining modality-specific information in working memory, leveraging existing specialized neural circuits for the efficient temporary storage of detailed sensory input (Adam et al. 2022). Despite its intuitive appeal it remains controversial whether vWM relies on the same neural substrate as visual perception or not, with strong empirical evidence supporting both sides of the debate (see Zhao et al. 2022; Sheremata et al. 2023 for support, and Bettencourt and Xu 2016; Iamshchinina et al. 2021 for critique; see also Tardiff and Curtis 2025, for review).

Given that the interindividual differences in vWM capacity are quite stable over time (Hockey and Geffen 2004; Johnson et al. 2013), it follows that there may be stable interindividual differences in brain architectonics underlying these behavioral effects. (Note that we use the terms ‘architectonics’ and ‘architectural’ to encompass both micro- and macrostructural tissue properties.) This offers a unique opportunity to use variability in vWM performance to uncover its neural origins via a correlational approach (Cronbach 1957; Hedge et al. 2018). Focusing on aspects of behavior that exhibit meaningful variance among individuals may overcome certain caveats of the experimental designs – where the focus is on differences between conditions, not among individuals – by reducing the likelihood of confounding neural correlates of vWM with other task-related processes. A number of studies have employed this interindividual-differences approach to identify the neural basis of vWM, albeit with varied results.

On the one hand, initial evidence demonstrated a link between vWM performance and V1 macrostructure (Bergmann et al. 2016). Thirty-one participants were tested on a delayed Gabor patch orientation discrimination task. Participants’ performance on this task was strongly linked to their individual V1 gray matter volume, with no similar correlations found in extrastriate visual cortex or higher-order areas. On the other hand, evidence exists that the gray matter volume within higher-order visual areas, such as the left lateral occipital region, is larger in individuals who retain a greater number of items in vWM (Machizawa, Driver, and Watanabe, 2020). Another study using a combination of two visual tasks – one focusing on spatial vWM and the other on object identity vWM of – showed no link with V1 macrostructure for either task (n=48). Instead, the spatial task was linked to gray matter volume in the inferior parietal lobules, whereas the object identity task was linked to gray matter volume in the insulae (Konstantinou et al. 2017). Moreover, a study of thirty-two participants on two typical visuospatial memory tasks revealed a strong link between total brain volume and behavioral performance. When this association with total brain volume was accounted for, local correlations in V1 and intraparietal sulcus were no longer significant (Dimond et al. 2019). Finally, a machine learning approach using a large multimodal MRI dataset (n=547) found widespread associations with vWM across the entire brain, with the strongest weights in motor and subcortical-cerebellum networks, whereas links within the early visual cortex were weak or absent (Xiao et al. 2021).

In summary, previous studies implicate various areas where gray matter volume seems to account for behavioral differences in vWM tasks, yet do not provide a clear answer regarding V1 involvement. The heterogeneity of these findings may be influenced by several factors. Below, we delineate two major contributors to this inconsistency and propose strategies to mitigate their impact.

The first source of noise arises from the reliance on macrostructural indices. Conventional structural MRI typically quantifies gray matter volume, cortical thickness, surface area, or curvature. While informative at the systems level, these metrics are indirect proxies for the underlying microstructural properties that ultimately determine neural computation. Macroscopic measures necessarily collapse across diverse cellular and molecular features – such as cyto- and myeloarchitecture – that are more proximally related to information processing.

The limitations of this approach are well illustrated by aging research. Aging is associated with microstructural remodeling, including demyelination (Callaghan et al., 2014) and increased tissue iron deposition (Ogg et al., 1998; Gracien et al., 2017). These alterations precede, and only gradually culminate in, detectable macroscopic changes in morphology (Seiler et al., 2021). By analogy, architectural properties of V1 that contribute to interindividual variability in vWM may manifest at the microstructural level without necessarily producing large-scale volumetric differences – particularly in healthy young adults. Moreover, macrostructural metrics are influenced by a broad constellation of tissue properties unrelated to vWM. Consequently, associations between macroscopic V1 measures and behaviour are likely to contain substantial variance orthogonal to the cognitive construct of interest, thereby attenuating effect sizes and contributing to inconsistent findings across studies.

Advances in quantitative MRI (qMRI) offer a principled means to address these limitations. Multiparametric qMRI enables in vivo assessment of biologically interpretable tissue parameters – indices sensitive to myelin content, iron concentration, or macromolecular composition – that may not be reflected in differences in volume or cortical thickness (Weiskopf et al., 2021). In contrast to conventional T1-weighted imaging, which yields scanner- and hardware-dependent mixed contrasts, qMRI techniques estimate quantitative tissue properties that are largely independent of acquisition hardware and therefore more comparable across sites (Weiskopf et al., 2013). These features enhance both reproducibility and interpretability. Importantly, because qMRI parameters correspond more directly to histological features, they facilitate integration with animal research. Accordingly, incorporating qMRI alongside conventional morphometric analyses provides a more comprehensive characterization of tissue properties underlying vWM, and reduces inherent confounds in purely macrostructural approaches.

A second source of noise concerns the behavioral aspects of brain–behavior association studies. For example, stimulus material is often limited within studies but varies widely across them. Whereas this is not necessarily a shortcoming (as WM is considered to be domain-specific and may differ, for example, between spatial vWM and object-based vWM; see Konstantinou et al. 2017), it makes it less likely that common neural markers are identified. A potentially more serious issue is that many brain-behavior correlation studies – including ROI-based studies – to date may have been underpowered (for a discussion, see Dubois and Adolphs 2016; Gratton et al. 2022; Marek et al. 2022; Rosenberg and Finn 2022). Depending on the stability and robustness of the behavioral effects, brain-wide association studies likely require upwards of a couple hundred participants to detect replicable findings (Kang et al. 2024). Many, if not all, behavioral effects include significant sources of noise because they are the product of many cognitive processes acting in unison. Therefore, to maximize the sensitivity, the relevance of any behavioral effect with respect to the actual latent construct of interest should be carefully considered. Ideally, the behavioral effect would not only be sensitive to the psychological constructs of interest but also distinguish between alternative hypotheses.

For example, if interindividual differences in vWM performance stem from architectural V1 differences, then vWM performance should exhibit consequences that mirror the known architectural properties of V1. The most striking architectural property of V1 is its retinotopy and the cortical magnification factor for foveal vision. Indeed, it is implicitly assumed that this is why V1 volume might be linked to vWM performance: more cortical real estate affords greater storage capacity (Jeong 2023). However, V1 architectonics are also marked by inhomogeneous representations of the visual field. As described in a review by Karim and Kojima (Karim and Kojima 2010), the visual field inhomogeneities include phenomena such as performance differences between the left and right or upper and lower visual fields, with the lower visual field dominating in tasks from contrast pattern recognition to motion perception.

Some authors have highlighted that the upper-versus-lower field effect in contrast sensitivity and acuity tasks is primarily driven by the vertical meridian, described as the vertical meridian asymmetry (VMA, Carrasco et al., 2001; Cameron et al., 2002). This is accompanied by an even stronger effect known as the horizontal-vertical anisotropy (HVA) – the dominance of performance along the horizontal compared to the vertical meridian. Those two phenomena are referred to as polar angle asymmetries (review, Himmelberg et al. 2023). While these asymmetries are already present in the organization of the photoreceptors and retinal-ganglion cells (Curcio et al. 1990; Curcio and Allen 1990; Song et al. 2011; Watson 2014), these retinal factors only partially explain the observed behavioral effects, pointing to the role of V1 (Kupers et al. 2019; Kupers et al. 2022), which manifests polar angle asymmetries both in both brain architecture Himmelberg et al., 202l; and function (Liu et al. 2006; O’Connell et al. 2016; Benson et al. 2021; Kurzawski et al. 2022; Kurzawski et al., 2025; Himmelberg et al 2022; Himmelberg et al. 2023; Himmelberg et al., 2025). In the following, we will use the term ‘visual field inhomogeneities’ when referring to both polar angle asymmetries and visual field asymmetries together.

If V1 is involved in vWM, it should be possible to detect these same asymmetries across the visual field and/or polar angles. Behavioral evidence indicates this is possible. A 2009 study demonstrated that both HVA and VMA extend to spatial frequency discrimination and perceived spatial frequency in visual short-term memory (Montaser-Kouhsari and Carrasco 2009). Visual field asymmetries in vWM performance have been also reported by Del Pin and colleagues (Del Pin et al. 2020), using an object-identification vWM task. Both identification accuracy and visibility ratings exhibited clear asymmetry patterns: the most prominent pattern closely resembled a typical HVA. An asymmetry was also present along the vertical axis but with an inverted direction – performance was on average better for the upper locations compared to the lower. These results suggest a potential involvement of V1 in vWM, although the observed pattern of visual field inhomogeneity –particularly the inverted VMA– diverges from canonical perceptual V1 asymmetries.

The first goal of the present study was to replicate the behavioral findings of Del Pin and colleagues (Del Pin et al. 2020) in a sample optimized for interindividual-differences research and explicitly test the robustness of the reported inverted VMA pattern. Critically, rather than relying solely on global performance metrics, we leveraged visual-field asymmetries to probe whether interindividual differences in performance systematically reflect the architectural idiosyncrasies of V1. This design increases construct specificity and provides a more stringent test of V1 involvement in vWM. The second goal was to overcome constraints associated with conventional morphometric approaches by combining established macrostructural measures of V1 with quantitative MRI–derived indices of underlying tissue properties. This allowed us not only to conduct a conceptual replication of Bergmann and colleagues (Bergmann et al., 2016), but also to extend their findings by testing whether specific microstructural characteristics of V1 account for interindividual differences in performance. By moving beyond purely volumetric descriptions, we aimed to identify biologically interpretable tissue parameters that may mechanistically link V1 architecture to behavior.

## Methods

### Participants

This study was conducted as part of the EU COST Action CA18106 (www.neuralarchcon.org) at the data-collection site in Kraków (Consciousness Lab, Jagiellonian University, Kraków, Poland). The broader consortium project includes an extensive behavioral test battery, MRI, EEG, and DNA sampling of neurotypical participants. The current study focuses exclusively on one visual working memory task from the larger behavioral battery and links it to interindividual variation specifically in the neuroarchitectural characteristics.

A total of 292 participants (185 F and 107 M), aged 18 to 40.9 years (mean = 23.9; SD = 4.47), took part in the study. The sample size was powered to account for the multiple comparisons correction required for whole-brain analyses. (For details see Section 1.3.2 of the Technical Annex of the Action; COST 2018). At the time of assessment, participants were physically healthy, reported normal or corrected-to-normal vision, and had normal hearing. Exclusion criteria included a history of brain damage or surgery, known neurological abnormalities, and the use of neuropharmacological or other medication that might affect neural states. Additionally, participants were excluded if they were pregnant, had contraindicating skin conditions, or a body habitus that precluded an MR scan. All participants met standard MRI safety criteria and signed written informed consent forms prior to participation. The project complies with the Declaration of Helsinki and was approved by the Ethics Committee of the Institute of Psychology, Jagiellonian University.

Participants attended two separate sessions. During the first session, they underwent a multiparametric mapping (MPM; Weiskopf, 2013) MRI scan (among other MRI sequences). During the second session, they performed the visual working memory task alongside other behavioral tasks. The two sessions were separated by an average of 11 days (SD = 6.6, range = 2 to 40).

### Behavioral task

The visual working memory task was similar to the object recognition task developed by Del Pin and colleagues (Del Pin et al. 2020). The task was created using the Psychopy Toolbox (Peirce et al. 2019), and comprised 256 trials. Each trial followed a temporal sequence: A fixation cross, (Thaler et al. 2013), designed to optimize gaze stability, was presented throughout the trial, excluding the response interval. Next, eight object images, each enclosed within a black square outline, appeared in a circular array centered at 6.8**°** eccentricity. After 1,000 ms, the object images disappeared, leaving the black squares to serve as location placeholders until the response screen. A 100 ms after stimulus offset, a cue – a black line pointing to one of the 8 locations – was then presented for 500 ms. Finally, after a 1,400 ms maintenance period, a probe object image appeared in the center of the screen. Participants indicated whether the probe was identical to the object previously presented in the cued location or not. A schematic representation of the trial structure is presented in **Figure 1A**.

**Figure 1.**
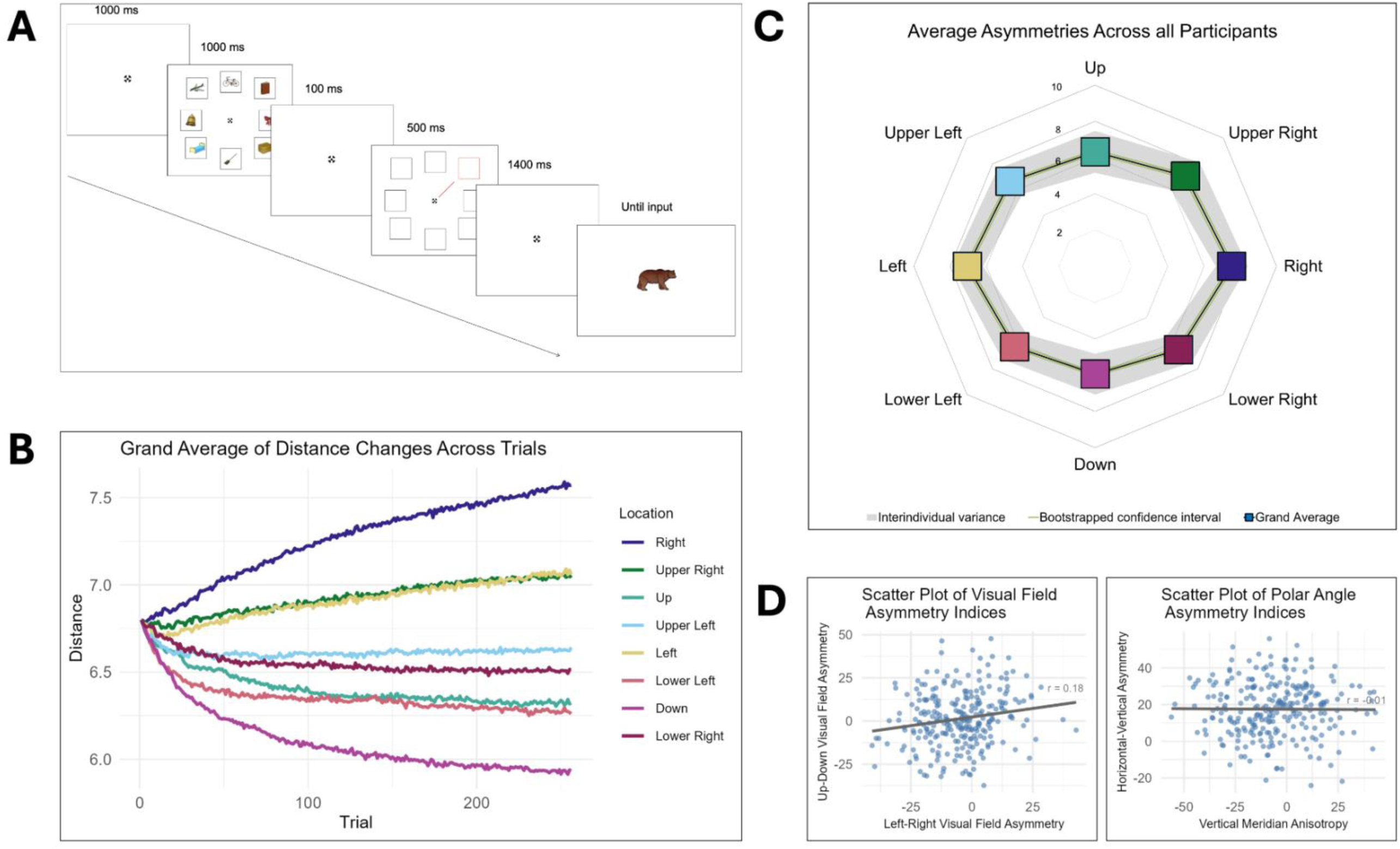
**(A)** A schematic representation of the trial structure of the visual working memory task used in the study. A set of eight targets was randomly chosen for each trial from a set of 200 images. The task included 256 trials, and the locations of the stimuli changed positions on a trial-by-trial basis. **(B)** Grand average of distance changes per location across trials for all participants (N = 262). The y-axis indicates the mean distance of the target locations from the center of the screen averaged across participants. Note, that distance is described in degrees of polar angle, and that all locations were initially presented at 6.8 degrees from the center of the screen; **(C)** Mean distance of each stimulus location from the center of the screen across participants (N = 262), calculated from the average distances in the final five trials of the experiment. Colored squares represent the group mean for each location. The grey shaded region indicates ±1 standard deviation across participants (interindividual variance), and the green line represents the 95% confidence interval of the bootstrapped mean (1000 bootstrap resamples). **(D)** Scatter plots illustrating relationships between the asymmetry indices. The correlation between the horizontal-vertical anisotropy and vertical meridian asymmetry correlation is insignificant (r = −0.01, p = 0.9), consistent with previous studies, whereas the correlation between upper–lower visual field asymmetry and left–right visual field asymmetry was weak but significant (r = 0.25, p < 0.01).

A set of 200 images (approximately 60% depicting man-made and 40% natural objects; no two images depicting the same object) was selected from the Snodgrass and Vanderwart object repository (Rossion and Pourtois 2004). Representative stimuli are available in **SI1**. The stimuli were chosen based on the ease of naming the depicted objects in Polish to ensure unambiguous recognition. The objects were scaled to a similar size with respect to the enclosing squares, maintaining clear gaps between the image and the square borders. Low-level properties (e.g., color, orientation, and spatial frequency) were not explicitly controlled.

The squares themselves extended 3.3×3.3 degrees of visual angle and were arranged at polar angles of 0**°**, 45**°**, 90**°**, 135**°**, 180**°**, 225**°**, 270**°**, and 315**°**. Most importantly, the distance of each square from fixation was adaptively and independently adjusted for each of the eight locations throughout the task using the QUEST staircase procedure (Watson and Pelli 1983; implemented in PsychoPy, Peirce et al. 2019). Each staircase adaptively targeted the stimulus eccentricity that yielded approximately 63% correct performance above chance (corresponding to an overall accuracy of 82%), consistent with bothPsychoPy QuestHandler documentation and the performance observed in our prior work (Del Pin et al. 2020).

The procedure assumed a standard Weibull psychometric function (50% guess rate, slope ≈ 3.5, lapse rate ≈ 0.01) and used an inverse mapping where the eccentricity was calculated as Distance (degrees) = 15 - QUEST_Intensity. Consequently, incorrect responses moved stimuli closer to fixation (decreasing task difficulty), whereas correct responses moved stimuli further away (increasing difficulty). The operational distance range was 3 to 15 degrees of visual angle, with all locations starting at 6.8°. Final distances for each location thus served as a proxy for performance, with greater distances indicating superior vWM ability at that visual field coordinate. **Figure 1B** depicts the average progression of the staircase for each of the eight locations.

The cued location and target image for each trial were determined at the beginning of the experiment. Each location was cued in 32 trials, and each of the 200 images appeared at a cued location at least once, with a random selection of 56 images appearing twice (256 trials total). The order of cued locations and target images was independently randomized for each participant. On every trial, seven distractor images were randomly chosen without replacement from the remaining set (excluding the target image) and assigned to the non-cued locations.

Whereas images varied in visual and semantic properties, the large stimulus pool (200 unique items) ensured that image-specific effects did not systematically influence the overall asymmetry patterns across the eight locations. Even though the images varied in their discriminability (as per post-hoc comparisons of natural and manmade objects, for example; see **SI3**), these differences should only influence the likelihood of responding correctly or incorrectly on a single trial, rather than disproportionately biasing specific location staircases over consecutive trials. Details regarding the individual images, their average discrimination rates, and comparisons between natural and manmade objects can be found in **SI3**.

The task was performed in a dimly lit room on a monitor driven by an NVIDIA GeForce RTX 2070 SUPER graphics card with a 60 Hz refresh rate and 1920 × 1080 screen resolution. Participants were seated approximately 55 cm from the screen and received written instructions. The experimenter verbally reminded participants to always maintain fixation at the center of the screen and avoid eye movements to the target locations. Responses were recorded via a standard keyboard using the left and right arrow keys.

### Behavioral analysis methods

Because the primary outcome of the vWM task was the final layout of the eight target locations at the end of the experiment, most participants were included in the analysis. We excluded trials with reaction times faster than 100 ms from the response screen onset or slower than 3 SD above the individual’s mean were removed. Participants were excluded if they met any of the following criteria: (1) had fewer than 205 clean trials remaining (80% of the experiment); (2) had an average performance level lower than 60% correct; (3) only used one of the response buttons (left/right) in more than 80% of the trials; or (4) did not finish the experiment within the allotted time. As a result, analyses were performed on 262 participants (170 F and 92 M; mean age = 23.77; SD = 4.26; 90.45% of the total sample). Note that for the brain-behavior analyses, 5 additional participants were removed because they had poor-quality MPM data from the MRI session (see next section).

The final position of the eight target locations was used to assess asymmetries in our vWM task. The final position was defined as the average distance of each location over the last five trials of the experiment to increase the robustness of the estimates. Other choices, such as averaging the distance over only the last five trials where a specific location was cued, led to very similar results (not reported). The distances of the target locations were subsequently aggregated into four summary indices of asymmetry and one summary index of overall vWM performance for each participant.

For clarity, we denote the final distance at each location using subscripts:

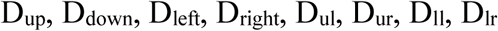

corresponding to the *up*, *down*, *left*, *right*, *upper-left*, *upper-right*, *lower-left*, and *lower-right* target locations, respectively. For example, D_up_ is the final distance of the item at the 12 o’clock location, D_right_ is the distance at the 3 o’clock location, D_down_ is the distance at the 6 o’clock location, and D_left_ is the final distance of the item at the 9 o’clock location.

In order to gain a complete overview of asymmetry effects, both polar angle and visual field asymmetries were analyzed. We used the horizontal-vertical anisotropy (HVA) and vertical meridian asymmetry (VMA) as indices of individual polar angle asymmetries. The indices were calculated based on established formulas (Himmelberg et al. 2022):

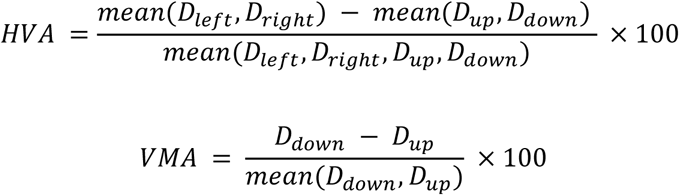

In addition to the polar angle asymmetries, broader visual field asymmetries were analyzed to explore upper versus lower visual field effect and left versus right visual field effects. To compare performance in the upper versus lower visual field, the distances of the *upper left, up,* and *upper right* locations were averaged and compared with analogous values for the *lower left, down,* and *lower right* locations. A similar approach was used to compare performance in the left (*upper left, left,* and *lower left* locations) vs. right (*upper right, right,* and *lower right* locations) visual field. The exact formulas for the visual field asymmetry indices were:

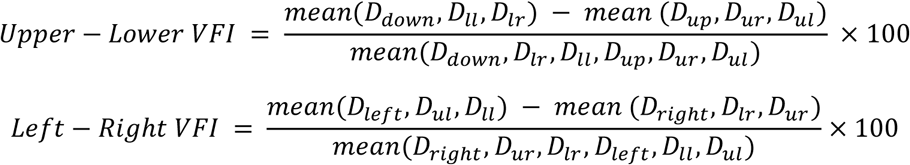

We specifically included locations on the visual meridians for the main analyses because perceptual performance asymmetries decrease away from the cardinal meridians and are no longer present at intercardinal locations (Carrasco et al. 2001; Corbett and Carrasco 2011; Abrams et al. 2012; Barbot et al. 2021), Nevertheless, meridian asymmetries and field asymmetries led to very similar results in this vWM task (see **SI4).**

Finally, we used average distance as a measure of overall vWM performance. This index was calculated by averaging the final distances across all eight target locations. This allowed us to test whether overall performance influenced asymmetry indices and whether potential neuroimaging links relate to vWM performance asymmetries. Furthermore, we use this index as an approximation of individual vWM performance limits in an attempt to conceptually replicate the findings of Bergmann and colleagues (Bergmann et al. 2016).

Given prior developmental findings showing changes in VMA across childhood and adolescence (Carrasco et al., 2023, 2025), as well as the significant correlation between individuals’ VMA and height (Carrasco et al., 2025), we additionally examined whether participant height was associated with asymmetry indices as supplementary control tests (see: **SI3**). The asymmetry indices were analyzed separately and compared to each other, or to other control variables such as eye dominance or height, via paired t-tests, Kruskal-Wallis rank tests, and Pearson correlations. P-values were adjusted using the Holm–Bonferroni correction for multiple comparisons. In cases where we aimed to determine whether two effects differed, we directly tested the difference between the corresponding effect estimates rather than inferring differences from their individual significance levels (Nieuwenhuis et al., 2011). For t-tests, this was implemented by conducting paired t-tests on the difference scores. Similarly, differences between correlation coefficients were formally compared using a bootstrap resampling approach (999 samples) with the *boot* package in R (Davison and Hinkley 1997; Canty and Ripley 2022). This allowed us to derive 95% confidence intervals for the observed differences.

All statistical analyses were performed using R version 4.3.1 (R Core Team 2023) and various packages, including *tidyverse* (Wickham et al. 2019), *effsize* (Torchiano & others, 2020), and *reshape* (Wickham 2007).

### MRI acquisition

The MRI data were acquired on a Siemens Magnetom Skyra 3T MR scanner. Each participant underwent resting-state fMRI, quantitative multi-parameter mapping (MPM), and diffusion-weighted imaging in a single scanning session lasting approximately one hour. This study uses only the MPM data. MPM allows for the acquisition of quantitative MRI (Weiskopf et al. 2013), with each contrast being sensitive to different microstructural aspects of neural tissue: magnetization transfer (MT) reflecting myelination (Bjarnason et al. 2005), effective proton density (PD) corresponding to water content (Lorio et al. 2019), longitudinal relaxation rate (R1) corresponding to the water mobility in its microenvironment and iron content (Callaghan et al. 2015), and effective transverse relaxation rate (R2*) representing iron accumulation (Draganski et al. 2011).

The MPM protocol was implemented based on recommended (Siemens) vendor sequences (Leutritz et al. 2020). Three-dimensional data acquisition consisted of three multi-echo spoiled gradient echo scans (i.e., fast low angle shot [FLASH] sequences with MT, T1, and PD contrast weighting). Additional reference radio-frequency (RF) scans were acquired. The acquisition protocol had the following parameters: TR for PDw and T1w contrasts: 18 ms; TR for MTw contrast: 37 ms; minimum/maximum TE for PDw, T1w, and MTw contrasts: 2.46/14.76 ms; flip angles for MTw, PDw, and T1w contrasts: 6°, 4°, and 25°, respectively; six equidistant echoes; 1 mm isotropic voxel size; field of view 224 × 256 ×176 mm; AP phase encoding direction; and a GRAPPA parallel imaging speedup factor of 2; acquisition times for T1w, PDw and MT were 3:50, 3:50, 7:52 min, respectively.

The acquisition of low-resolution three-dimensional spoiled gradient echo volumes was executed using both the RF head coil and the body coil. This dual acquisition facilitated the generation of a relative net RF receive field sensitivity (B1−) map for the head coil. This approach achieved rapid acquisition by maintaining a low isotropic spatial resolution of 4 mm³, a short echo time (TE) of approximately 2 ms, and a reduced flip angle of 6°, without parallel imaging acceleration or partial Fourier encoding. This procedure of capturing volume pairs with the head and body coils was repeated before acquiring each of the MT, PD, and T1 contrasts.

### MRI preprocessing

The raw MPM data were converted into four quantitative maps (MT, PD, R1, R2*) via the recommended map creation module of the hMRI-toolbox, a dedicated SPM add-on for MPM data analysis (Tabelow et al. 2019). Map creation includes an auto-reorientation step and two steps correcting quantitative MRI estimates for spatial receive and transmit field inhomogeneities, respectively. At this stage, all resulting maps were visually inspected; pronounced movement artifacts or abnormal brain anatomy (i.e., large ventricles or visible tumors) led to the exclusion of 5 additional participants (a total of 257 participants included in the MRI analysis). Further preprocessing steps followed the voxel-based quantification (VBQ) pipeline, which parallels the standard voxel-based morphometry (VBM) pipeline for volumetric macrostructural analysis in SPM. The pipeline was performed on individual MT maps first, with the resulting transformations subsequently applied to the remaining maps. In short, the VBQ pipeline first segments each image into gray matter, white matter, and cerebrospinal fluid. Next, all individual images are spatially registered to a group average template using the DARTEL algorithm. Finally, each tissue type is smoothed with a 4 mm FWHM kernel. Note that the final step deviates slightly from the standard VBM pipeline. For MPM data, a specialized tissue-weighted smoothing algorithm is applied to avoid partial volume effects at tissue boundaries that would distort the quantitative estimates.

To perform surface-based macrostructural analyses in addition to the volumetric analysis described above, a synthetic T1w image was generated from the R1 and PD maps. First, both maps were thresholded to resemble more closely the typical statistics of a T1-weighted image. This is necessary because freesurfer has been optimized specifically for T1w images and expects a certain range of input (Fischl et al., 1999; Lutti et al., 2014). Using FSL commands, the R1 map was divided by itself twice, thresholded at zero, and multiplied by 1,000. The PD map was thresholded at zero and multiplied by 100. Subsequently, the mri_synthesize FreeSurfer command was applied to create a synthetic FLASH image based on the R1 and PD maps. The argument for optimal gray and white matter contrast weighting was used with the parameters TR=20, TE=30, and flip angle=2.5. Finally, the synthetic T1w image was scaled (divided by four) to comply with FreeSurfer’s expected intensity range.

Surface reconstructions for estimating cortical thickness were generated using the standard pipeline in CAT12 toolbox (Gaser et al. 2024). The pipeline uses a projection-based thickness method (Dahnke et al. 2013) and surface refinement to model the cortical sheet and folding patterns. Topological defects are automatically repaired using spherical harmonics, and individual central surfaces are spatially registered via spherical mapping with minimal distortions. Finally, local thickness values were transferred to the Freesurfer ‘fsaverage’ template.

### MRI analysis

Statistical analyses were conducted for both volumetric MPM data and surface-based cortical thickness data. VBQ was performed using the SPM12 toolbox (Wellcome Trust Centre for Neuroimaging 2014), and cortical thickness analysis was performed on synthetic T1 maps using the CAT12 toolbox (Gaser et al. 2024).

### ROI analysis

For VBQ, the region of interest (ROI) analysis focused on the primary visual cortex V1 to investigate the sensory recruitment hypothesis. In addition to V1, we tested two non-sensory regions known for their role in visual working memory processing: the intraparietal sulci (IPS; Bettencourt and Xu 2016; Iamshchinina et al. 2021), and the frontal eye fields (FEF; Offen et al. 2010; Jerde et al. 2012; Lefco et al. 2020). Importantly, the FEF include the superior precentral sulcus (sPCS; Bedini et al. 2023), an area linked with the maintenance of vWM representations (Jerde et al. 2012).

Analysis was performed using atlas-based masks created with the WFU Pick Atlas toolbox (Maldjian et al. 2003). Individual 3D masks covered Brodmann area (BA) 17 and BA 6 (representing V1 and FEF bilaterally, respectively). Additional masks of the early visual cortex were created for control analyses to assess the systematicity of V1-related results. Selected regions included left and right BA17; dorsal and ventral BA17 (analyzed both bilaterally and separately for each hemisphere); BA18 (corresponding to bilateral V2), and combined BA17+18. All masks created with the WFU Pick Atlas toolbox were dilated by 1 voxel. ROIs were subdivided into dorsal and ventral banks using the median MNI z-coordinate of each mask. The division occurred at z = 4 mm (median z of the right, left, and bilateral BA 17 masks).

Spherical masks of IPS were created with the MarsBaR toolbox (Brett et al. 2002). The MNI coordinates (x,y,z) were 38, −42, 44 for the right IPS and −40, −42, 46 for the left IPS, with a radius of 10 mm. Coordinates were identified using the Neurosynth database by searching the term “intraparietal sulcus” in the meta-analyses section. We selected the peak coordinates that were most consistently reported across studies; these overlapped with regions identified in a separate search for the keyword “working memory” and were within 6 mm of coordinates used in several working memory studies (e.g., Knops et al., 2006; Mayer et al., 2007; Schneiders et al., 2011; Lu et al., 2016; Gayet et al., 2017). Inspection of the probabilistic visual topography atlas (Wang et al., 2015) indicates that these spheres partially overlap retinotopically organized IPS4-5 and lie in close proximity to IPS3, suggesting the ROI corresponds to the posterior-middle visuotopically organized intraparietal cortex.

For all indices, analysis was performed as follows: All voxels within the chosen ROI were averaged for each participant. Then, two linear mixed-effect models (LMMs) were fitted and compared using a likelihood ratio test (LRT). The null model included total intracranial volume (TIV), age, and sex as nuisance covariates, and participant as a random effect. The alternative model comprised the asymmetry index as a predictor, the same nuisance covariates, and participant as a random effect. This analysis was performed for each of the four MPM contrasts (MT, PD, R1, and R2*), each ROI, and each behavioral index separately. Results were corrected for multiple comparisons within each index using the False Discovery Rate (FDR) method. As a supplementary control analysis, the volumetric models were re-estimated including participant height as a covariate (see **SI6**).

The ROIs for cortical thickness analysis were derived from the Human Connectome Project multimodal parcellation (HCP-MMP1.0) atlas (Glasser et al. 2016). The regions of interest were analogous to those for VBQ and included V1, IPS1, and FEF. The statistical procedure paralleled the VBQ analysis: the basic model included age and sex as nuisance covariates, with the behavioral index added as a predictor to the full model. The contribution of each index to cortical thickness was assessed using LRTs comparing the full and reduced models.

### Whole brain analysis

Whole-brain analyses were conducted using the previously-estimated gray matter models. Data were analyzed separately for HVA, VMA, upper-lower and left-right visual field indices, and average distance of all target locations. LRTs compared models including the behavioral index, sex, age, and TIV as predictors, against null models without the behavioral indices. If the behavioral indices significantly impact the brain architectonics, voxel clusters should emerge. The Holm-Bonferroni method was used to control for the family-wise error rate (FWER).

## Results

### Behavioral results

The behavioral task in this study was designed to map inhomogeneities in vWM performance across the visual field. We summarized these spatial inhomogeneities via polar angle and visual field asymmetry indices and tested whether their distributions differed statistically significantly from zero. The horizontal-vertical anisotropy (HVA) reached an average value of 17.5 (min = −24.14, max = 55.88, SD = 14.7) and the distribution differed significantly from zero (t(261) = 19.26, p <0.001, Cohen’s d = 1.19). This indicates that, on average, performance was better along the horizontal axis compared to the vertical axis, as expected. The vertical meridian asymmetry (VMA) reached an average of −6.15 (min = −56.02, max = 43.18, SD = 19.64), and this distribution also differed significantly from zero (t(261)= −5.07, p <0.001, Cohen’s d = −0.31).

Regarding visual field asymmetries, the left-right asymmetry reached an average value of −4.83 (min = −40.93, max = 42.86, SD =13.98) and the distribution differed significantly from zero (t(261)= −5.59, p <0.001, Cohen’s d = −0.35). The superior performance in the right visual field was not related to eye-dominance (χ2 = 0.07, p = 0.94). The upper-lower asymmetry reached an average value of −11.43 (min = −51.72, max = 23.16, SD = 15.29), and its distribution differed significantly from zero (t(261)= −12.1, p<0.001, Cohen’s d = −0.75). Similar to VMA, the polarity of the upper-lower asymmetry indicates better performance in the upper compared to the lower visual hemifield, on average. This similarity was further supported by a significant correlation between the two indices (r(260)=0.58, p<0.001).

To summarize, the results demonstrate clear spatial inhomogeneities in a vWM task both along the polar angles and across the visual field. We next examined whether interindividual differences in polar angle asymmetries or visual field asymmetries were related to each other. A Pearson’s correlation between VMA and HVA yielded a null result (r(260) = −0.01, p = 0.9, 95% CI [–0.13, 0.11]). The upper-lower and left-right asymmetries, on the other hand, were significantly correlated (r(260) = 0.25, p<0.01, 95% CI [0.13, 0.36]). A test of these two correlations confirmed that the latter was systematically stronger than the former (95% bootstrap CI: [-0.39, −0.1]) indicating that visual field asymmetries share a source of variance whereas polar angle asymmetries do not (see Himmelberg et al. 2022).

The pattern of results mostly confirmed our a priori expectations, including the inverted VMA direction. In an effort to eliminate potential confounding sources, we tested whether interindividual variability in VMA could be linked to participant height (as the viewing angle of the screen certainly differed between tall and short participants). This correlation was not significant (r(260) = −0.02, p=0.79). The variability of VMA was also not related to eye-dominance (χ2 = 1.04, p = 0.31; See **SI3** for correlations with other indices). We therefore conclude that, contrary to many visual perception tasks, performance in our vWM task is better along the upper than the lower vertical meridian.

Finally, we assessed the average distance of the eight target locations at the end of the task. As the staircase was used to adjust the distance from the center, the final average distance was highly correlated with accuracy (r = 0.86, p < 0.001). Therefore, we used average distance as a measure of overall vWM performance. The average distance varied between 5.39° and 8.08° (mean = 6.67, SD = 0.52). Note that polar angle asymmetry patterns were not confounded by average distance; VMA was not significantly correlated with average distance (r = −0.01, p = 0.83), and HVA manifested a weak correlation (r = 0.14, p = 0.03). Conversely, visual field indices were moderately correlated with average distance (r = 0.45, p < 0.001 for up-down and r = 0.48, p <0.001 for left-right). This effect was expected, as visual field asymmetries rely on the average distances. Additionally, we assessed whether average reaction time was linked with accuracy or average distance. These correlations were negligible and not statistically significant (see **SI5**).

### Brain-behavior relationships

The behavioral results confirm systematic inhomogeneities in vWM performance across the visual field. These inhomogeneities were statistically significant for both polar angle and visual field asymmetry indices. We next investigate whether these behavioral patterns are linked to interindividual differences in brain architecture.

Given that the hypotheses of this work specifically target the early visual cortex and its potential links to working memory performance, we focus primarily on ROI analyses of V1 (and V2; see **SI7**), with the IPS and FEF as control regions. The latter two regions have been consistently linked to vWM processing, but do not necessarily provide detailed perceptual representations of memory content. Therefore, if interindividual differences in behavior are linked to architectural differences in the IPS/FEF but not V1, our data would speak against the sensory recruitment hypothesis. If, on the other hand, behavioral patterns are linked to V1 architecture (regardless of IPS/FEF results) we would interpret this as supporting the sensory recruitment hypothesis. Whole-brain analyses are presented for completeness.

In the following sections, we first analyze volumetric differences across the four MPM maps, where each map reflects different microstructural tissue properties. Second, interindividual differences in cortical thickness – derived from the synthetic T1w image – are related to the behavioral results.

### Volumetric results

A set of regression analyses relating the behavioral asymmetry indices to the four MPM maps across the three ROIs yielded null results for the IPS and FEF (For details, see **Table 1**). However, three MPM parameters – R2*, PD, and R1 – showed significant associations with the behavioral VMA index in V1. Specifically, the R2* map is primarily indicative of iron content in brain tissue, while PD and R1 maps are linked to total water content and a combination of molecular water mobility and iron content. This indicates that participants with a more pronounced behavioral VMA tended to have denser neural tissue in V1, characterized by lower water content and higher iron concentration compared to participants with less asymmetry along the vertical meridian (see **Figure 2**).

**Figure 2.**
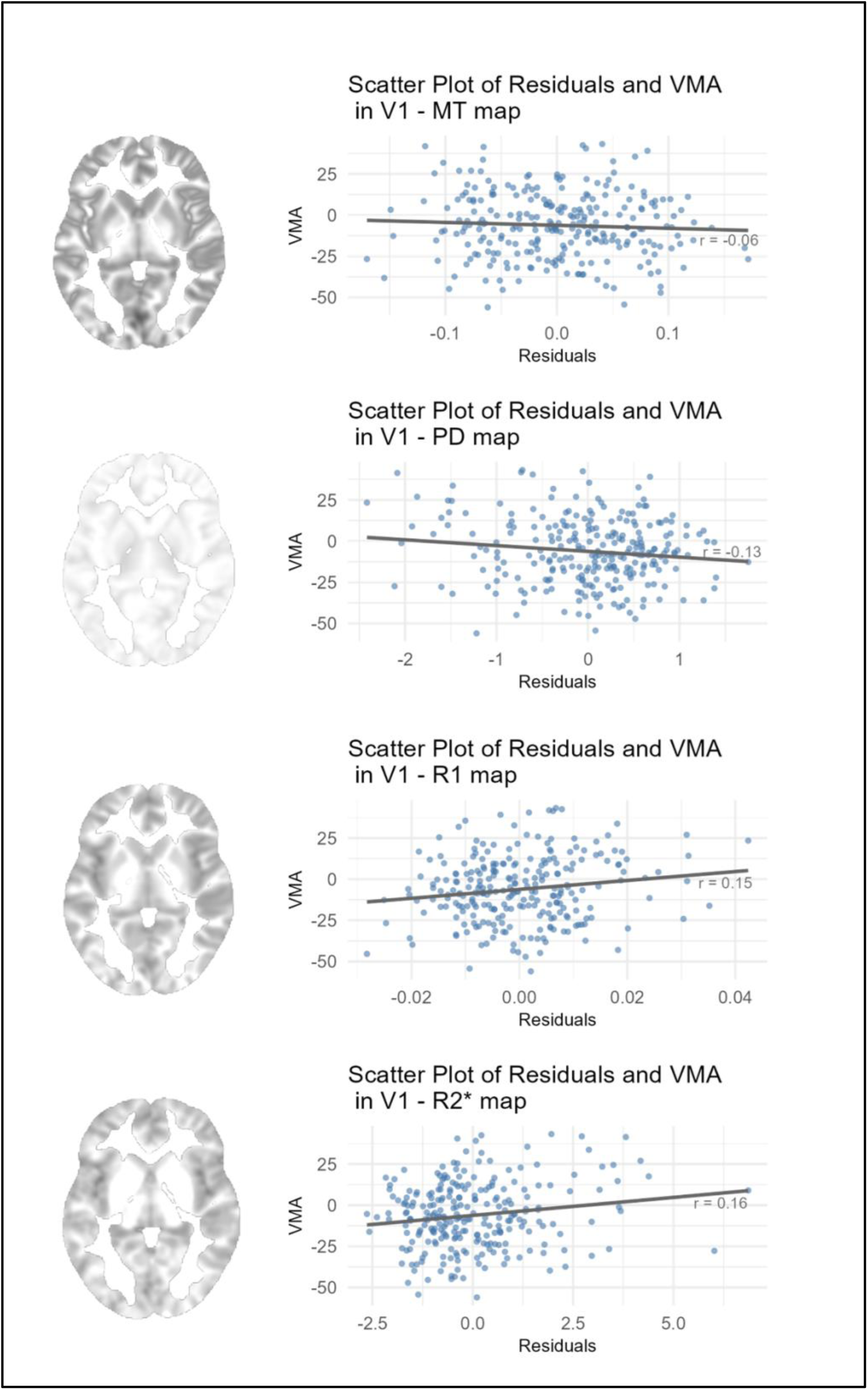
Scatter plots showing the relationship between the VMA index and model residuals from linear mixed-effects models predicting MPM values within V1 for the four quantitative maps (MT, PD, R1, and R2*). Models included sex, age, and total intracranial volume as covariates, with participant included as a random effect. Each data point corresponds to one participant (N = 257). Each panel displays Pearson correlation coefficient (*r*) between the residual MPM values and the VMA. Likelihood ratio tests comparing models with and without the VMA predictor revealed significant associations for PD (χ²(1)= 4.4, p=0.048), R1 (χ²(1)=6.08, p=0.027) and R2* (χ²(1)=6.76, p=0.027), whereas MT (χ²(1)= 0.85, p=0.36) did not reach significance (see Table 1 for full statistical results). Anatomical images illustrate example slices of the respective MPM quantitative maps.

**Table 1.** Volumetric and cortical thickness results. Results from the Likelihood Ratio Tests (LRTs) compare complex models (df = 7) against null models (df = 6) to explain ROI-wise variance. P-values were corrected for different MPM contrasts within each index using the False Discovery Rate (FDR) method. Cortical thickness results display F-statistics and uncorrected p-values for each index and ROI.

|  | Bilateral V1 multiparametrical maps |  |  |  |  |  |  |  | V1 cortical thickness |  |  |  |
| --- | --- | --- | --- | --- | --- | --- | --- | --- | --- | --- | --- | --- |
|  | MT |  | PD |  | R1 |  | R2* |  | Left V1 |  | Right V1 |  |
| Index | $\chi^2(1)$ | p | $\chi^2(1)$ | p | $\chi^2(1)$ | p | $\chi^2(1)$ | p | F | p | F | p |
| Horizontal-Vertical Anisotropy | 0.45 | 0.729 | 0.39 | 0.729 | 0.12 | 0.729 | 0.13 | 0.729 | 1.06 | 0.305 | 0.244 | 0.622 |
| Vertical Meridian Asymmetry | 0.85 | 0.36 | 4.4 | 0.048 | 6.08 | 0.027 | 6.76 | 0.027 | 0.61 | 0.435 | 1.6 | 0.207 |
| Upper-Lower Visual Field Index | 1.95 | 0.172 | 1.86 | 0.172 | 1.97 | 0.172 | 0.71 | 0.172 | 0.61 | 0.435 | 0.45 | 0.504 |
| Left-Right Visual Field Index | 0.2 | 0.65 | 1.69 | 0.39 | 3.4 | 0.26 | 0.9 | 0.46 | 0.8 | 0.373 | 0.01 | 0.933 |
| Average Distance | 0.93 | 0.867 | <0.001 | 0.984 | 0.04 | 0.984 | 0.64 | 0.867 | 0.29 | 0.589 | 0.1 | 0.754 |
|  | Bilateral IPS multiparametrical maps |  |  |  |  |  |  |  | IPS cortical thickness |  |  |  |
|  | MT |  | PD |  | R1 |  | R2* |  | Left IPS |  | Right IPS |  |
| Index | $\chi^2(1)$ | p | $\chi^2(1)$ | p | $\chi^2(1)$ | p | $\chi^2(1)$ | p | F | p | F | p |
| Horizontal-Vertical Anisotropy | 0.05 | 0.82 | 0.51 | 0.82 | 1.29 | 0.82 | 0.21 | 0.82 | 0.18 | 0.676 | 2.56 | 0.111 |

**Table 1. Volumetric and cortical thickness results.**
|  |  |  |  |  |  |  |  |  |  |  |  |  |
| --- | --- | --- | --- | --- | --- | --- | --- | --- | --- | --- | --- | --- |
| Vertical Meridian Asymmetry | 0.004 | 0.95 | 0.09 | 0.95 | 0.61 | 0.87 | 0.68 | 0.87 | 0.69 | 0.406 | 1.36 | 0.245 |
| Upper-Lower Visual Field Index | 0.28 | 0.802 | 0.22 | 0.802 | 0.06 | 0.802 | 0.79 | 0.802 | 0.001 | 0.970 | 1.74 | 0.188 |
| Left-Right Visual Field Index | 3.94 | 0.19 | 0.85 | 0.85 | 0.71 | 0.4 | 0.27 | 0.61 | 0.93 | 0.335 | 0.19 | 0.661 |
| Average Distance | 2.68 | 0.388 | 0.13 | 0.72 | 1.69 | 0.388 | 0.25 | 0.72 | 2.35 | 0.127 | 2.97 | 0.08 |
|  | Bilateral FEF multiparametrical maps |  |  |  |  |  |  |  | FEF cortical thickness |  |  |  |
|  | MT |  | PD |  | R1 |  | R2* |  | Left FEF |  | Right FEF |  |
| Index | $\chi^2(1)$ | p | $\chi^2(1)$ | p | $\chi^2(1)$ | p | $\chi^2(1)$ | p | F | p | F | p |
| Horizontal-Vertical Anisotropy | <0.001 | 0.997 | 1.07 | 0.603 | 0.009 | 0.997 | 1.2 | 0.603 | 0.47 | 0.495 | 0.51 | 0.475 |
| Vertical Meridian Asymmetry | 0.92 | 0.777 | 0.08 | 0.777 | 1.24 | 0.531 | 1.33 | 0.531 | 0.16 | 0.686 | 2.66 | 0.104 |
| Upper-Lower Visual Field Index | 0.92 | 0.45 | 2.49 | 0.45 | 0.15 | 0.695 | 1.35 | 0.45 | 0.55 | 0.458 | 2.55 | 0.111 |
| Left-Right Visual Field Index | 0.2 | 0.707 | 0.14 | 0.707 | 0.34 | 0.707 | 0.19 | 0.707 | 0.16 | 0.688 | 0.2 | 0.656 |
| Average Distance | 0.13 | 0.746 | 0.34 | 0.746 | 1.3 | 0.746 | 0.11 | 0.746 | 0.83 | 0.362 | 0.21 | 0.646 |

Additional analyses revealed that these associations between the PD, R1, and R2* maps and the behavioral VMA were only robust in left V1, particularly in its ventral subdivision, and were not observed in right V1. These effects were not consistent when analyzing V2 or the combined V1+V2 ROI (see **SI7**). The same analyses for upper-lower visual field asymmetry did not reach significance. Nevertheless, direct comparisons between the PD, R1, and R2* correlation strengths revealed that the effects with the second-highest χ2 value within each MPM map were not systematically weaker than the VMA effects with the highest χ2 value (95% bootstrap CIs: [-0.1855, 0.1294] for PD, [-0.1514, 0.2278] for R1, [-0.0719, 0.2345] for R2*). This indicates that the current analysis may lack the power to uncover smaller effects present in other indices or ROIs.

To further examine whether behavioral asymmetries were reflected in corresponding architectural asymmetries within V1, we computed volumetric asymmetry indices based on dorsal–ventral and left–right subdivisions of BA17. This accounted for the retinotopic organization of the early visual cortex and explored the spatial specificity of the observed effects. These analyses did not reveal significant associations between architectural and behavioral asymmetry indices (see **SI8**).

In addition to the behavioral asymmetry indices, identical analyses were performed for average distance to assess any architectural correlates of overall vWM performance. These analyses aimed to conceptually replicate the findings of Bergmann and colleagues (Bergmann et al. 2016), who reported a link between V1 gray matter volume and individual performance limits in a vWM task. However, none of the regression models including average distance as a predictor, were significant in our dataset.

Finally, we conducted a whole-brain analysis. No clusters were found for any of the asymmetry indices in any of the MPM maps.

### Cortical thickness results

Cortical thickness was analyzed similarly to the volumetric MPM data; specifically, the average cortical thickness within the three ROIs was regressed against behavioral indices and nuisance covariates. None of the asymmetry indices or the average distance were significant predictors in these models (for details, see **Table 1**).

In an exploratory whole-brain analysis (implemented in CAT12), we tested cortical thickness across all atlas-defined regions using Holm-Bonferroni correction for multiple comparisons. This analysis revealed significant correlations between left-right visual field asymmetry and cortical thickness in two regions: left V3 (F = 9.87, p = 0.041) and right Brodmann area 25 (BA 25, subgenual anterior cingulate cortex; F = 4.45, p = 0.036). Based on these exploratory findings, we performed post-hoc examinations of V3 and BA 25 for the remaining behavioral indices. This was characterized by largely null results, with the exception of a significant association in V3 for average distance (F = 8.81, p = 0.003). The whole-brain analysis yielded no significant results for the other visual field indices.

## Discussion

This study examined whether architectural properties of the early visual cortex explain asymmetries in visual working memory (vWM) performance. Behavioral results of the vWM task are consistent with well-established perceptual asymmetries, including a strong horizontal–vertical anisotropy (HVA, Montaser-Kouhsari and Carrasco 2009), a left–right asymmetry favoring the right hemifield (Karim and Kojima 2010). We also observed an inverted vertical meridian asymmetry (VMA), with better performance in the upper than the lower vertical meridian of the visual field, contrasting with some prior reports (Montaser-Kouhsari and Carrasco 2009). Individual asymmetry indices were then tested as predictors of cortical micro- and macrostructure. While no effects emerged in IPS or FEF, V1 architecture was linked to the VMA across the PD, R1, and R2* maps. Additional subdivision analysis (see **SI8**) revealed that this effect was primarily driven by the ventral portion of left V1, corresponding approximately to the visual field region with the highest behavioral performance in our task. Because individual retinotopic mapping was not available, the atlas-based subdivisions of V1 should be considered approximations, and the spatial specificity of these effects should be interpreted with caution.

Importantly, we identified a connection between the cortical thickness of left V3 and both left-right visual asymmetry and the average distance index. As task performance was generally better in the right, upper right, and lower right locations, the association within the left visual cortex – which represents the right visual field (Sereno et al. 1995) – is anatomically meaningful. V3 has been linked with conscious perception and stimulus discrimination (Salminen-Vaparanta et al. 2019), which could account for the enhanced performance in the right hemifield among participants with a thicker V3. Macrostructural differences in extrastriate regions have also been linked to individual differences in vWM performance; for example, larger gray matter volume in the left lateral occipital cortex has been associated with greater VWM capacity (Machizawa, Driver, and Watanabe, 2020).

Furthermore, V3 is part of the early visual cortex and shares many architectural properties with the striate cortex. Given our reliance on atlas-based definitions rather than direct retinotopic mapping, findings across V1-V3 should be interpreted broadly within the context of the early visual cortex. In this context, we regard the V3 finding as consistent with the sensory recruitment hypothesis. Additionally, we observed a relationship between left-right asymmetry and the thickness of the right subgenual area (BA 25). Since this region is primarily involved in emotional regulation (Mayberg et al. 1999) and autonomic functions (Sudheimer et al. 2015; Wallis et al. 2017), we consider this finding incidental.

The behavioral results yielded several noteworthy patterns. First, performance in the right visual hemifield was better than in the left, which was unanticipated. Numerous studies did not exhibit this effect, due to the bilateral field advantage, both hemispheres should have separate attentional resources (Zhang et al. 2017; Strong and Alvarez 2020), allowing for unified performance across hemifields. However, the right visual hemifield has occasionally manifested improved performance in cognitive tasks (Karim and Kojima 2010). While specialized recognition for tools in the right visual field has been reported (Garcea et al. 2012), stimulus type (natural or manmade, including tools) did not affect performance in our study (see: **SI2**).

Moreover, the upper-lower visual field asymmetries were inverted relative to previous literature, which describes the lower visual field dominance in various perceptual and cognitive tasks, both regarding the whole lower hemifield (Karim and Kojima 2010) and the VMA (Carrasco et al. 2001; Corbett and Carrasco 2011; Abrams et al. 2012; Himmelberg et al., 2023). However, some studies suggest that humans recognize objects more accurately in the upper than in the lower visual field (Chambers et al. 1996; Zito et al. 2016; Del Pin et al. 2020), while others proved increased working memory performance in the lower visual hemifield (Genzano et al. 2001), particularly the lower vertical meridian (Montaser-Kouhsari and Carrasco 2009). Given these conflicting reports, further studies are needed to explore the origins of the inverted VMA observed here.

Average distance served as a marker of overall vWM performance. It correlated moderately with upper-lower and left-right hemifield asymmetries, but its link with HVA was negligible, and with the VMA, non-significant. This may point to the unique properties of the VMA, which is represented cortically along the longitudinal fissure (Glickstein and Whitteridge 1987).

Horizontal-vertical anisotropy, the most pronounced asymmetry pattern in our task, was not significantly linked to V1 architectonics in any MPM maps. In contrast, the comparatively small and inverted VMA was linked with individual differences in V1 architectonics. This finding diverges from existing literature linking both HVA and VMA to the early visual cortex (Himmelberg et al. 2023). Nevertheless, because HVA and VMA are known to be uncorrelated (Himmelberg et al. 2020; Barbot et al. 2021; Purokayastha et al. 2021; Himmelberg et al. 2022), our findings also suggest they arise from distinct neural underpinnings. This aligns with the finding that polar angle asymmetries are at least partially heritable (Benson et al. 2021).

We found the VMA to be associated with three MPM parameters: PD, R1 and R2*. The PD contrast primarily reflects free water content (Lorio et al. 2019) whereas R1 is sensitive to water mobility and iron content of the neural tissue (Callaghan et al. 2015). The R2* map reflects the iron accumulation (Draganski et al. 2011). Importantly, these quantitative MRI measures provide information about cortical microstructure that extends beyond the anatomical size of the macrostructures.

This distinction is relevant because prior studies have focused primarily on macrostructural or functional measures (e.g., Bergmann et al., 2016; Dimond et al., 2019; Zhao et al., 2022; Rademaker et al., 2019). Quantitative MRI adds a new dimension by capturing variability in tissue composition, informing not only *where* in the brain macrostructural variability relates to behavior, but also *what* properties of the tissue may contribute to these relationships. Our findings suggest that interindividual differences in spatial vWM asymmetries may not be explained solely by macrostructural properties of V1, such as volume or cortical thickness, but also by differences in its underlying microstructural composition.

Within the sensory recruitment hypothesis, this is particularly informative. If vWM maintenance relies on neural encoding in the early visual cortex, then variability in architectural properties x, beyond its overall size, may contribute to performance differences. MPM measures complement macrostructural metrics and refine our understanding of how anatomical variability relates to cognitive variability.

Previous work has linked iron-sensitive MPM parameters to variability in cognitive performance and aging-related cognitive decline (e.g., Raven et al., 2013; Spence et al., 2020; Granziera et al., 2015). Similar markers have been associated with factors such as age, smoking (Hagemeier et al. 2015), and high body mass index (Pirpamer et al. 2016), known risk factors for later cognitive impairment (Waisman et al., 2016; Qu et al., 2020). However, those studies have not examined microstructural variability specifically within the early visual cortex. The present findings extend this line of research by linking quantitative MRI markers of V1 microstructure to spatial asymmetries in vWM performance.

While MT and PD maps manifest high reliability, the reliability of R1 and R2* maps can be more limited in healthy young adults, with high intraclass correlation coefficient (ICC) for MT (0.88) and PD (0.87), and moderate for R1 (0.73) and R2* (0.78; Wenger et al., 2022). This could be linked to the generally low variability of iron deposition and molecular water mobility in this age group. That study included a smaller sample (15 participants) and did not include the early visual cortex as a region of interest for the reliability examination. However, our findings in a larger sample (N=256) suggest that these markers are sensitive to behavioral variance. Future studies across a broader age range are needed to establish how increased iron accumulation in V1 impacts asymmetries in vWM performance.

Our findings regarding V1 surface differences are in partial agreement with Bergmann and colleagues (Bergmann et al. 2016): we found a link between vWM performance and the architectonics of V1 (and V3, which is also part of the early visual cortex). Importantly, our findings yielded smaller effect sizes and were specific to certain quantitative maps and asymmetry indices. Unlike Bergmann et al., we did not observe a link between V1 cortical thickness and vWM performance, although our V3 cortical thickness finding corroborates the role of early visual cortex in working memory.

Despite promising results, our study has several limitations. First, due to the absence of a chinrest and eye-tracking we cannot entirely rule out effects related to scanning bias or changes in viewing distance (Carrasco et al. 1995; Rezec and Dobkins 2004). However, stimuli were presented in random locations, and the fixation cross was designed specifically to stabilize fixation (Thaler et al. 2013). Additionally, post-hoc analyses found no impact of participant height (a proxy of viewing angle) or eye dominance on asymmetry patterns (see **SI3**). Finally, our study did not include individual retinotopic mapping, which would have improved the accuracy of V1 selection compared to the atlas-based approach and allowed for more granular structural analysis.

## Acknowledgements

We thank the entire NeuralArchCon consortium (www.neuralarchcon.org). We thank Katarzyna Hat and Monika Ostrogorska for collecting the MRI data. We thank Tomasz Kostka, Wiktoria Orłowska, and the C-lab interns between 2020-2022 for their assistance in the behavioral data collection.

