## Supplementary material for "Visual Field Inhomogeneities and the Architectonics of Early Visual Cortex Shape Visual Working Memory": SI1, SI2, SI3, SI4, SI5, SI6, SI7, SI8

Supplementary materials

1. Experimental stimuli

The stimuli for the task were chosen from the Snodgrass and Vanderwart *Like* object repository (Rossion and Pourtois 2004);  for details, see the methods section). A randomly chosen subset of experimental stimuli is presented in Figure 1.

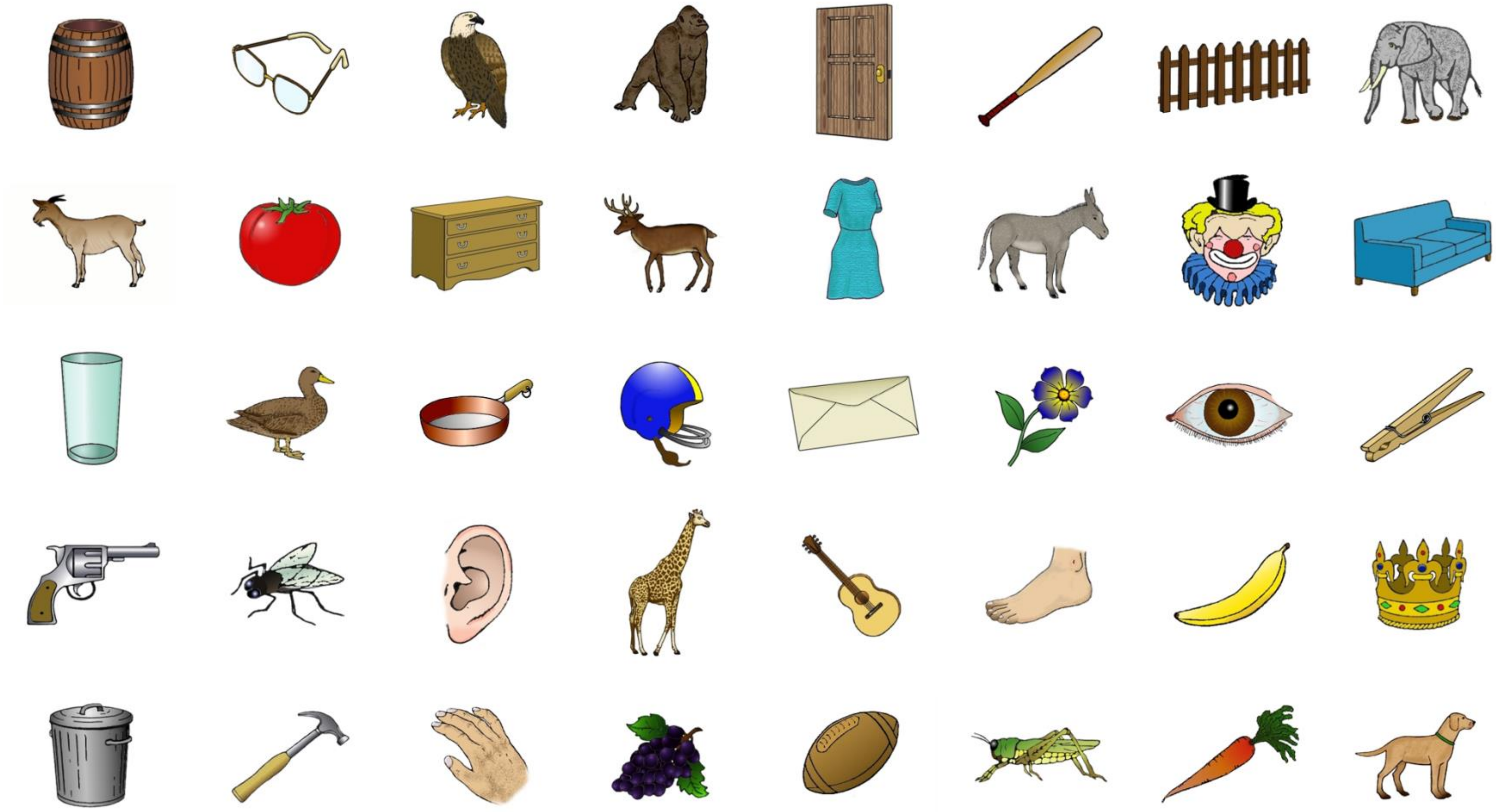

Figure 1. A randomly chosen subset of experimental stimuli.

2. Stimuli type effect on performance

A paired t-test was performed to check whether objects from one category (manmade or natural) were perceived with higher accuracy. The results show that responses to natural stimuli were more often correct than responses to manmade stimuli ($t = 3.8$, $p < 0.001$, Cohen’s $d = 0.54$, $df = 198$), with average accuracy rate reaching 0.74 for natural, and 0.71 for manmade objects ($SD = 0.05$ and 0.04, respectively). The tendency for higher accuracy for natural stimuli was consistent across locations (see Figure 2B), yet any specific participants did not carry the results; as presented on Figure 2C, participants who tended to have higher accuracy in natural stimuli tended to have a high accuracy for the manmade stimuli as well. This was confirmed by Pearson's correlation ($r(260) = 0.57$, $p < 0.001$). The average accuracy for each stimulus used in the procedure are presented in Figure 2A.

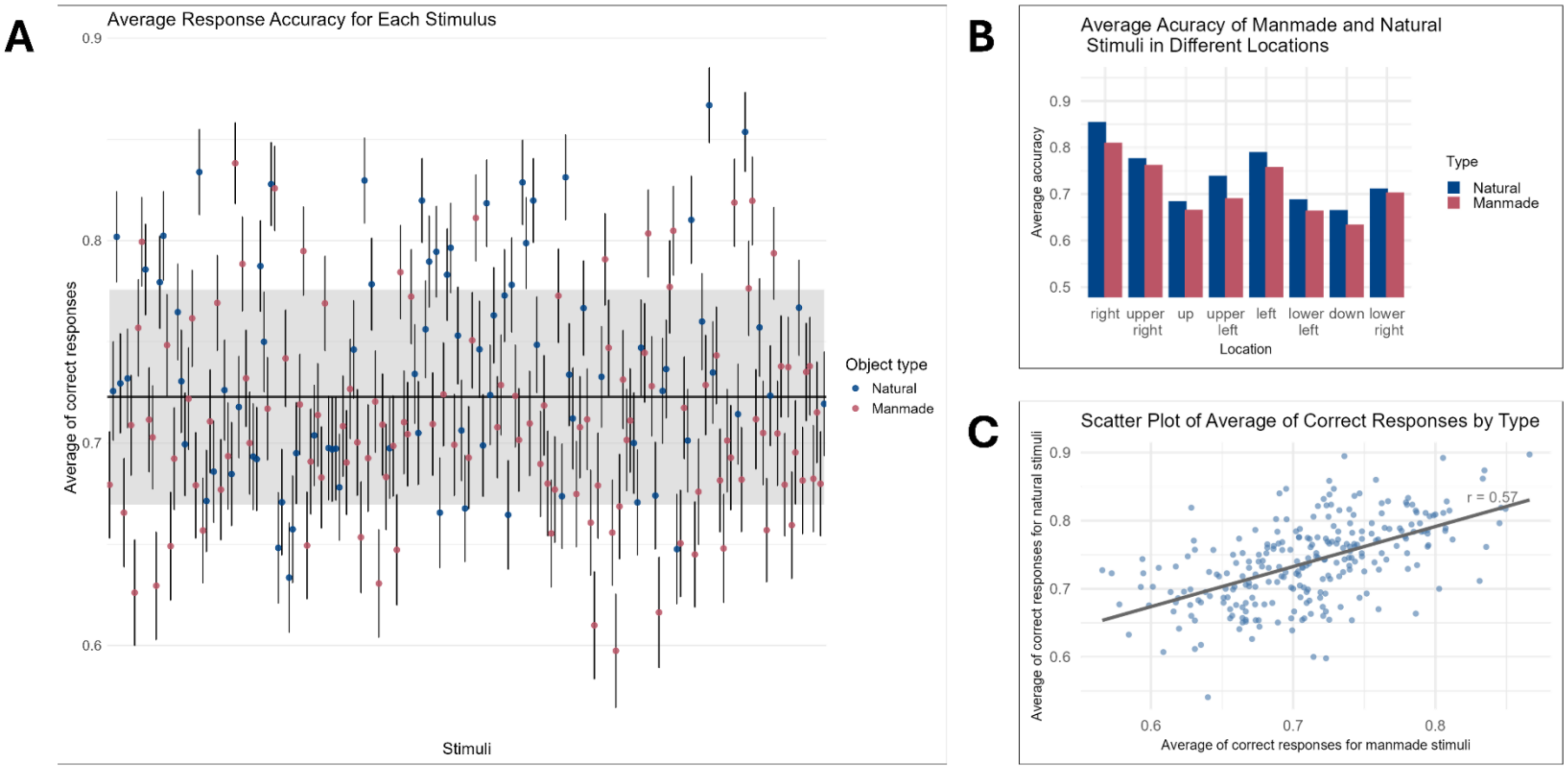

A
Average Response Accuracy for Each Stimulus
Average of correct responses
Stimuli
Object type
Natural
Manmade
B
Average Acuracy of Manmade and Natural Stimuli in Different Locations
Average accuracy
right
upper right
up
upper left
left
lower left
down
lower right
Location
Type
Natural
Manmade
C
Scatter Plot of Average of Correct Responses by Type
Average of correct responses for natural stimuli
Average of correct responses for manmade stimuli
r = 0.57

*Figure 2*. Average response accuracy for each stimulus used for the procedure. The naturally occurring objects are marked in blue, and the manmade in red. The thin black lines represent the standard error for the average response accuracy for each stimulus across participants. The thick black line represents the average accuracy across all objects, and the gray shading – the confidence interval (M = 0.71, SD = 0.05)

3. Eye dominance and height

The effects of eye dominance and height on the asymmetry indices were tested in order to assess whether they could confound the data. The reasoning behind testing the link between the height on the indices was that the vertical meridian anisotropy is known to vary between children and adults, (Carrasco et al. 2022) appearing and developing throughout adolescence (Carrasco et al. 2023; Carrasco et al., 2025), which is believed to be linked to the development of the preference for perceiving the lower half of the visual field, as this part of the visual field tends to hold relevant information on a day-to-day basis (Carrasco et al., 2023). For completeness, we performed Pearson's correlation tests for the effect of height on all asymmetries, yet the results were null, consistent with previous research (Carrasco et al., 2023; see: Table 1).

The effect of eye-dominance on the asymmetry indices was tested post-hoc after the data analysis showed that our participants had higher accuracy for recognizing objects appearing in the right visual field. However, the Kruskal-Wallis tests yielded null results (see Table 2).

| | **r(260)** | **t** | **p-value (uncorrected)** | **p-value (corrected)** |
|---|---|---|---|---|
| Horizontal-Vertical Anisotropy | 0.14 | 2.26 | 0.03 | 0.82 |
| Vertical Meridian Asymmetry | -0.02 | -0.27 | 0.79 | 0.82 |
| Upper-lower VFI | 0.01 | 0.23 | 0.82 | 0.82 |
| Left-right VFI | -0.02 | -0.26 | 0.79 | 0.82 |

*Table 1. Pearson's correlation between individual asymmetry indices and height of the participants.*

| | $\chi2$ (1) | p-value (uncorrected) | p-value (corrected) |
|---|---|---|---|
| Horizontal-Vertical Anisotropy | 0.36 | 0.55 | 0.79 |
| Vertical Meridian Asymmetry | 1.04 | 0.31 | 0.79 |
| Upper-lower VFI | 0.14 | 0.7 | 0.79 |
| Left-right VFI | 0.07 | 0.79 | 0.79 |

*Table 2.. Kruskall-Wallis test results of the effect of eye dominance on individual indices.*

4. On- and non-meridian asymmetries

The visual field analysis of the on-meridian upper versus lower visual field (*up* vs *down*) demonstrated a small effect size (t = 7.5, df = 261, Cohen's d = 0.6, p<0.001), while the non-meridan upper versus lower visual field analysis (mean(*upper right, upper left*) vs mean(*lower right, lower left*)) showed a smaller t value, but a medium effect size (t = 6.88, df = 261, Cohen's d = 0.56, p<0.001). Both isocentric (*left* vs *right*) and non isocentric (mean(*lower left, upper left*) vs mean(*lower right, lower right*)) left versus right visual field analyses showed a small effect size. (for on-axis: t = -747, df = 261, Cohen's d = 0.-0.53, p<0.001; for off-axis: t = 5.52, df = 261, Cohen's d = 0.42, p<0.001). The summary of comparisons of different visual field effects can be found in Table 3.

| | **t(261)** | **p uncorrected** | **p corrected** | **Cohen's d** |
|---|---|---|---|---|
| Upper-lower VF vs vertical axis | -0.79 | 0.42 | 0.43 | -0.04 |
| Left-right VF vs horizontal axis | 1.58 | 0.12 | 0.17 | 0.0 |
| HVA vs VMA | 8.13 | <0.001 | <0.001 | 0.71 |
| Left-right VF vs upper-lower VF | -10.35 | <0.001 | <0.001 | -0.93 |
| Left-right VFI vs non-meridian left-right VFI | -1.58 | 0.12 | 0.17 | -0.06 |
| Upper-lower VFI vs non-meridians upper-lower VFI | -0.79 | 0.43 | 0.43 | -0.02 |

*Table 3*. Formal comparisons of the differences. The comparisons were made by calculating the t-tests of the differences within the indices. FDR correction was applied to account for the multiple comparisons.

5. Accuracy, mean distance, and reaction time

Pearson's correlation coefficients were calculated in order to test whether there is a link between the accuracy of the performance in the task and the average reaction time, as well as between the reaction time and the mean distance of all locations from the center throughout the last 5 trials. Both results did not yield statistical significance (r(260) = -0.12, p = 0.052 and p(26) = -0.1, p = 0.11 for the average reaction time and mean distance and accuracy and average reaction time respectively). For details see Figure 3.

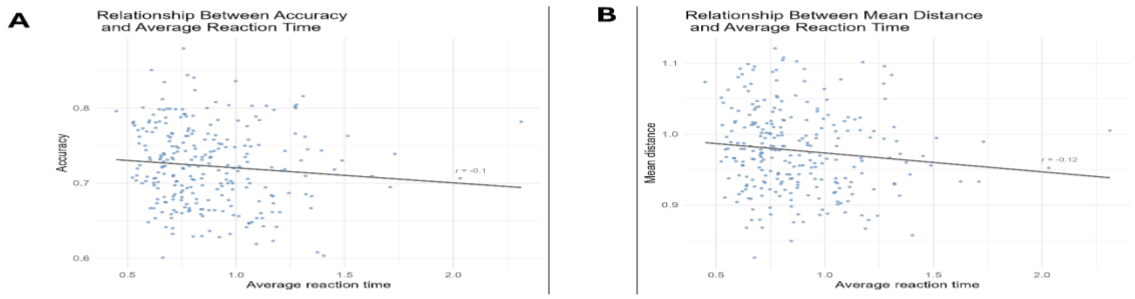


*Figure 3. A: scatter plot of accuracy in the task performance and the average reaction time of each participant. B: Scatter plot of the mean distance and average reaction time of each participant.*

6. Brain-behavior analyses including height as covariate

In line with the reasoning described in SI3, we cosidered if height should be added as a covariate into the volumetric models. To assess that, we ran a Pearson's corelation coeffitient matrix to test whether any of used covariates share substantial variance. The results show moderate correlation of height with both sex and total intracranial volume (for details see Table 3). Taken together with the null correlations of height and behavioral indices, height was excluded as a covariate in the main brain analysis.

As a post-hoc robustness check, we re-estimated the main volumetric model for bilateral V1 and the VMA index including participant height as an additional covariate. The results are presented in Table 4. The overall pattern of results remained consistent with the primary analyses.

| | Sex | Age | Total Intracranial Volume | Height |
|---|---|---|---|---|
| Sex | 1 | -0.0765 | 0.5883 | 0.5573 |
| Age | -0.0765 | 1 | 0.0135 | 0.0310 |
| Total Intracranial Volume | 0.5883 | 0.0135 | 1 | 0.5032 |
| Height | 0.5573 | 0.0310 | 0.5032 | 1 |

*Table 3.* Pearson's correlation coefficient matrix of covariates included in brain-behavior models.

| | **Bilateral V1 multiparametrical maps** | | | | | | | |
|---|---|---|---|---|---|---|---|---|
| | **MT** | | **PD** | | **R1** | | **R2*** | |
| **Index** | χ2(1) | p | χ2(1) | p | χ2(1) | p | χ2(1) | p |
| Vertical Meridian Anisotropy (height included as a covariate) | 0.83 | 0.36 | 4.24 | 0.052 | 6.12 | 0.027 | 6.67 | 0.027 |

*Table 4.* Results of the Likelihood Ratio tests comparing more complex models (df = 7) explaining the variance of the ROI voxels against simpler ones (df = 6). The p-values were corrected for comparisons of different maps within the index using the FDR

7. Regions of interest

To further account for the retinotopic organization of V1 and to examine the spatial specificity of the observed effects, we repeated the primary likelihood ratio model comparisons within anatomically defined subdivisions of BA17. Specifically, BA17 was subdivided into left and right hemispheres to approximate contralateral visual hemifield representations, and into dorsal and ventral portions to approximate lower and upper visual field representations, respectively. In addition, quadrant-level subdivisions (left dorsal, left ventral, right dorsal, right ventral) were examined to further explore spatial specificity.

All analyses followed the same nested model comparison procedure described in the Methods section. For completeness, control analyses in BA18 (V2) and in a combined BA17+18 mask are also included in Table 4.

Across these subdivisions, significant effects were limited to the association between vertical meridian anisotropy and R2* within BA17. The observed association showed spatial specificity, being primarily localized to the ventral portion of left BA17. Effects in other subdivisions did not reach statistical significance

| | **Left V1 multiparametrical maps** | | | | | | | |
|---|---|---|---|---|---|---|---|---|
| | **MT** | | **PD** | | **R1** | | **R2*** | |
| **Index** | χ2(1) | p | χ2(1) | p | χ2(1) | p | χ2(1) | p |
| Horizontal-Vertical Anisotropy | 0.94 | 0.953 | 0.83 | 0.953 | 0.38 | 0.953 | 0.03 | 0.953 |
| Vertical Meridian Asymmetry | 2.36 | 0.125 | **4.82** | **0.038** | **5.76** | **0.033** | **7.13** | **0.03** |
| Upper-Lower Visual Field Index | 3.63 | 0.227 | 1.78 | 0.244 | 1.16 | 0.282 | 2.3 | 0.243 |
| Left-Right Visual Field Index | 1.43 | 0.23 | 3.38 | 0.13 | 4.18 | 0.13 | 2.36 | 0.17 |
| Average Distance | 0.04 | 0.989 | 0.23 | 0.989 | 0.3 | 0.989 | <0.001 | 0.989 |
| | **Right V1 multiparametrical maps** | | | | | | | |
| | **MT** | | **PD** | | **R1** | | **R2*** | |
| **Index** | χ2(1) | p | χ2(1) | p | χ2(1) | p | χ2(1) | p |
| Horizontal-Vertical Anisotropy | 0.01 | 0.952 | 0.93 | 0.952 | 0.004 | 0.952 | 0.63 | 0.952 |
| Vertical Meridian Asymmetry | 0.06 | 0.814 | 2.68 | 0.137 | 4.82 | 0.113 | 2.67 | 0.137 |
| Upper-Lower Visual Field Index | 0.03 | 0.87 | 1.34 | 0.493 | 2.2 | 0.493 | 0.05 | 0.870 |
| Left-Right Visual Field Index | 0.27 | 0.89 | 0.18 | 0.89 | 1.63 | 0.81 | 0.02 | 0.9 |
| Average Distance | 1.63 | 0.404 | 0.36 | 0.404 | 0.06 | 0.404 | 2.4 | 0.404 |
| | **Dorsal V1 multiparametrical maps** | | | | | | | |
| | **MT** | | **PD** | | **R1** | | **R2*** | |
| **Index** | χ2(1) | p | χ2(1) | p | χ2(1) | p | χ2(1) | p |
| Horizontal-Vertical Anisotropy | < 0.001 | 0.994 | 0.94 | 0.899 | 0.14 | 0.949 | 0.57 | 0.899 |

| | | | | | | | | |
|---|---|---|---|---|---|---|---|---|
| Vertical Meridian Asymmetry | 0.015 | 0.904 | 0.51 | 0.633 | 2.79 | 0.189 | 4.11 | 0.170 |
| Upper-Lower Visual Field Index | 1.82 | 0.355 | 2.18 | 0.355 | 0.77 | 0.46 | 0.55 | 0.46 |
| Left-Right Visual Field Index | 0.009 | 0.922 | 1.52 | 0.433 | 1.65 | 0.433 | 0.10 | 0.922 |
| Average Distance | 0.89 | 0.677 | 1.02 | 0.677 | 0.36 | 0.677 | 0.677 | 0.677 |
| | **Ventral V1 multiparametrical maps** | | | | | | | |
| | **MT** | | **PD** | | **R1** | | **R2*** | |
| **Index** | χ2(1) | p | χ2(1) | p | χ2(1) | p | χ2(1) | p |
| Horizontal-Vertical Anisotropy | 1.19 | 0.831 | 0.16 | 0.831 | 0.41 | 0.831 | 0.05 | 0.831 |
| Vertical Meridian Asymmetry | 2.12 | 0.145 | **5.24** | **0.029** | **6.15** | **0.026** | **6.34** | **0.026** |
| Upper-Lower Visual Field Index | 1.53 | 0.372 | 1.17 | 0.372 | 2.00 | 0.372 | 0.62 | 0.431 |
| Left-Right Visual Field Index | 0.52 | 0.472 | 1.16 | 0.392 | 3.16 | 0.301 | 1.10 | 0.392 |
| Average Distance | 0.65 | 0.836 | 0.24 | 0.836 | 0.003 | 0.953 | 0.73 | 0.836 |
| | **Right Dorsal V1 multiparametrical maps** | | | | | | | |
| | **MT** | | **PD** | | **R1** | | **R2*** | |
| **Index** | χ2(1) | p | χ2(1) | p | χ2(1) | p | χ2(1) | p |
| Horizontal-Vertical Anisotropy | 0.13 | 0.723 | 1.43 | 0.464 | 0.57 | 0.599 | 1.45 | 0.464 |
| Vertical Meridian Asymmetry | 0.41 | 0.656 | 0.20 | 0.656 | 2.65 | 0.222 | 2.54 | 0.222 |
| Upper-Lower Visual Field Index | 0.60 | 0.585 | 1.15 | 0.585 | 0.80 | 0.585 | 0.009 | 0.924 |
| Left-Right Visual Field Index | 0.50 | 0.638 | 1.44 | 0.460 | 2.32 | 0.460 | 0.007 | 0.933 |
| Average Distance | 1.93 | 0.568 | 0.36 | 0.568 | 0.33 | 0.568 | 0.76 | 0.568 |
| | **Left Dorsal V1 multiparametrical maps** | | | | | | | |

| | MT | | PD | | R1 | | R2* | |
|---|---|---|---|---|---|---|---|---|
| **Index** | χ2(1) | p | χ2(1) | p | χ2(1) | p | χ2(1) | p |
| Horizontal-Vertical Anisotropy | 0.15 | 0.862 | 0.26 | 0.862 | 0.03 | 0.862 | 0.03 | 0.862 |
| Vertical Meridian Asymmetry | 0.38 | 0.539 | 1.34 | 0.330 | 2.36 | 0.249 | 4.36 | 0.148 |
| Upper-Lower Visual Field Index | 3.20 | 0.131 | 3.85 | 0.131 | 0.51 | 0.476 | 2.73 | 0.131 |
| Left-Right Visual Field Index | 0.79 | 0.497 | 1.42 | 0.497 | 0.46 | 0.497 | 0.60 | 0.497 |
| Average Distance | 0.007 | 0.934 | 1.96 | 0.647 | 0.29 | 0.890 | 0.18 | 0.890 |
| | **Right Ventral V1 multiparametrical maps** | | | | | | | |
| | **MT** | | **PD** | | **R1** | | **R2*** | |
| **Index** | χ2(1) | p | χ2(1) | p | χ2(1) | p | χ2(1) | p |
| Horizontal-Vertical Anisotropy | 0.02 | 0.922 | 0.48 | 0.922 | 0.13 | 0.922 | 0.01 | 0.922 |
| Vertical Meridian Anisotropy | 0.01 | 0.904 | 3.84 | 0.100 | 4.60 | 0.100 | 2.12 | 0.194 |
| Upper-Lower Visual Field Index | 0.03 | 0.865 | 0.99 | 0.637 | 2.33 | 0.507 | 0.15 | 0.865 |
| Left-Right Visual Field Index | 0.08 | 0.882 | 0.02 | 0.882 | 0.61 | 0.882 | 0.07 | 0.882 |
| Average Distance | 0.85 | 0.411 | 1.42 | 0.411 | 0.68 | 0.411 | 2.99 | 0.335 |
| | **Left Ventral V1 multiparametrical maps** | | | | | | | |
| | **MT** | | **PD** | | **R1** | | **R2*** | |
| **Index** | χ2(1) | p | χ2(1) | p | χ2(1) | p | χ2(1) | p |
| Horizontal-Vertical Anisotropy | 1.46 | 0.909 | 0.01 | 0.911 | 0.47 | 0.911 | 0.08 | 0.911 |
| Vertical Meridian Asymmetry | 3.11 | 0.078 | **4.85** | **0.037** | **5.64** | **0.035** | **6.86** | **0.035** |
| Upper-Lower Visual Field Index | 3.07 | 0.320 | 0.92 | 0.336 | 1.09 | 0.336 | 1.92 | 0.332 |

| | | | | | | | | |
|---|---|---|---|---|---|---|---|---|
| Left-Right Visual Field Index | 1.38 | 0.241 | 3.20 | 0.148 | 4.95 | 0.104 | 2.59 | 0.144 |
| Average Distance | 0.07 | 0.912 | 0.01 | 0.912 | 0.22 | 0.912 | 0.01 | 0.912 |
| | **V2 multiparametrical maps** | | | | | | | |
| | **MT** | | **PD** | | **R1** | | **R2*** | |
| **Index** | χ2(1) | p | χ2(1) | p | χ2(1) | p | χ2(1) | p |
| Horizontal-Vertical Anisotropy | 0.13 | 0.885 | 0.02 | 0.885 | 0.03 | 0.885 | 0.62 | 0.885 |
| Vertical Meridian Asymmetry | 1.72 | 0.189 | 1.99 | 0.189 | 4.4 | 0.071 | 5.63 | 0.07 |
| Upper-Lower Visual Field Index | 3.73 | 0.214 | 1.86 | 0.231 | 1.9 | 0.231 | 0.7 | 0.403 |
| Left-Right Visual Field Index | 0.05 | 0.85 | 1.08 | 0.6 | 2.84 | 0.37 | 0.04 | 0.85 |
| Average Distance | 0.48 | 0.604 | 0.27 | 0.604 | 0.44 | 0.604 | 0.57 | 0.604 |
| | **V1+V2 multiparametrical maps** | | | | | | | |
| | **MT** | | **PD** | | **R1** | | **R2*** | |
| **Index** | χ2(1) | p | χ2(1) | p | χ2(1) | p | χ2(1) | p |
| Horizontal-Vertical Anisotropy | 0.29 | 0.795 | 0.28 | 0.795 | 0.002 | 0.966 | 0.93 | 0.795 |
| Vertical Meridian Asymmetry | 0.95 | 0.406 | 0.69 | 0.406 | 3.09 | 0.157 | 5.3 | 0.085 |
| Upper-Lower Visual Field Index | 3.57 | 0.236 | 1.29 | 0.416 | 1.02 | 0.416 | 0.28 | 0.594 |
| Left-Right Visual Field Index | 0.006 | 0.93 | 0.59 | 0.89 | 2.46 | 0.47 | 0.02 | 0.93 |
| Average Distance | 0.55 | 0.587 | 0.3 | 0.587 | 0.33 | 0.587 | 0.76 | 0.587 |

*Table 5.* Results of the Likelihood Ratio tests comparing more complex models (df = 7) explaining the variance of the ROI voxels against simpler ones (df = 6). The p-values were corrected for comparisons of different maps within each index using the FDR

8. Volumetric asymmetry indices

For completeness, we conducted additional analyses to examine whether visual hemifield asymmetries in behavior were reflected in corresponding structural asymmetries within V1. To this end, structural asymmetry indices were computed analogously to the behavioral indices, using mean MPM values extracted per participant from anatomically defined V1 subdivisions. Specifically, we denote:

$$V_{dorsal}, V_{ventral}, V_{left}, V_{right}$$

corresponding to the average MPM values within dorsal BA17, ventral BA17, left, and right BA17, respectively. Structural asymmetry indices were computed as follows:

$$Upper - Lower\ Structural\ Asymmetry = \frac{V_{dorsal} - V_{ventral}}{mean(V_{dorsal}, V_{ventral})} \times 100$$

$$Left - Right\ Structural\ Asymmetry = \frac{V_{right} - V_{left}}{mean(V_{right}, V_{left})} \times 100$$

The order of subtraction was chosen to mirror the behavioral indices and account for V1 retinotopy: ventral V1 corresponds to the upper visual hemifield, dorsal V1 to the lower visual hemifield, and each hemisphere represents the contralateral visual field. Descriptive statistics of the structural asymmetry indices are reported in Table 6.

Structural asymmetry indices were computed separately for each MPM contrast. Nested model comparisons were then performed following the same procedure as in the primary volumetric analyses, with the structural asymmetry index used as the dependent variable. The null model included total intracranial volume, age, and sex as nuisance covariates, along with a random intercept for participant. The alternative model additionally included the corresponding behavioral asymmetry index as a predictor. Results were corrected for multiple comparisons within each index using the FDR procedure.

No significant associations were observed between behavioral asymmetry indices and structural asymmetry measures (see Table 7).

| | V1 - volumetric asymmetry descriptive statistics | | | | | | | | | | | | | | | |
|---|---|---|---|---|---|---|---|---|---|---|---|---|---|---|---|---|
| | **MT** | | | | **PD** | | | | **R1** | | | | **R2*** | | | |
| **Index** | Mean | SD | Min | Max | Mean | SD | Min | Max | Mean | SD | Min | Max | Mean | SD | Min | Max |
| Upper-Lower Visual Field Index | 14.35 | 9.37 | -1.99 | 47.08 | 4.30 | 3.28 | -3.51 | 16.91 | -4.34 | 4.71 | -21.87 | 3.94 | -41.43 | 11.62 | -84.83 | -14.13 |
| Left-Right Visual Field Index | -3.81 | 12.51 | -27.03 | 32.83 | -200.38 | 1.18 | -202.73 | -196.99 | -2.99 | 2.89 | -13.88 | 6.04 | 2.78 | 15.96 | -46.94 | 38.24 |

Table 6. Descriptive statistics of the volumetric asymmetry indices.

| | **V1 - volumetric asymmetry analysis** | | | | | | | |
|---|---|---|---|---|---|---|---|---|
| | **MT** | | **PD** | | **R1** | | **R2*** | |
| **Index** | $\chi2(1)$ | p | $\chi2(1)$ | p | $\chi2(1)$ | p | $\chi2(1)$ | p |
| Upper-Lower Visual Field Index | 0.34 | 0.566 | 0.46 | 0.566 | 1.28 | 0.566 | 0.33 | 0.566 |
| Left-Right Visual Field Index | 1.95 | 0.190 | 1.83 | 0.190 | 1.75 | 0.190 | 3.09 | 0.190 |

Table 7. Results of the Likelihood Ratio tests comparing more complex models (df = 7) explaining the variance of the ROI voxels against simpler ones (df = 6). The p values were corrected for comparisons of different maps within each index using the FDR.
